# Experience-Dependent Plasticity of Thalamoprefrontal Circuitry Characterizes Learning Across Species

**DOI:** 10.1101/2025.11.23.689976

**Authors:** Isabella S. Elkinbard, Sina Sadeghzadeh, Michelle Hedlund, Valerie Tsai, Theodore T. Ho, Karl Deisseroth, Vivek P. Buch

## Abstract

Learning is an adaptive process which is thought to require precise coordination among multiple brain regions over time. Thalamoprefrontal circuitry is central to multiple domains of cognitive control and executive function, yet its evolution during learning is not characterized. In this study, we explored thalamoprefrontal circuit signaling in mice and connectivity dynamics in humans over the course of learning a homologous psychomotor task. In mice, we targeted a genetically encoded Ca^2+^indicator to the mediodorsal-prefrontal (MdT-dmPFC) projection and used fiber photometry to record synaptic dorsomedial PFC (dmPFC) activity. As learning sessions progressed, all mice demonstrated a decrease in reaction time (RT) performance on successful trials (p<0.001). In concomitant synaptic MdT-dmPFC circuit activity across sessions, we observed a significant increase in activity during the anticipatory (p<0.01) and reward retrieval (p<0.01) periods, and a non-significant trend towards an increase in preparatory activity (p<0.15). However, following the learning period, during task re-exposure, we observed a significant shift in circuit activity, away from anticipatory (p<0.001) and towards the preparatory period (p<0.001) over the course of re-exposure sessions. Furthermore, we observed an emergence of a learner (decrease in RT) and a non-learner group (increase in RT) during the task re-exposure period. Over the course of a single analogous task session in humans, we also observed a learner and a non-learner group. When analyzing the thalamoprefrontal regional connectivity dynamics of early and late trials for learners, we observed a significant increase in low frequency and decrease in high frequency synchrony (connectivity tilt) in thalamoprefrontal pathways during preparatory and anticipatory periods. Interestingly, pairwise interactions specifically between the anterior corona radiata (ACR) and superior frontal gyrus (SFG), the human homolog to the genetically targeted MdT-dmPFC circuitry in mice, in fact demonstrated a robustly opposite connectivity tilt effect distinguishing learners from non-learners (Cohens *d* > 2). Overall, these findings may provide cross-species evidence of novel, conserved thalamoprefrontal circuit mechanisms of adaptive learning.

## Main

The brain has a remarkable ability to adapt and change its structure and function in response to external experiences and internal cues ^1^. Performing a single task can alter neuronal connectivity and signaling efficacy within specific circuits—an adaptive mechanism that enables the brain to adjust to an ever-changing environment, known as neuroplasticity^2,3,4,5^. While mechanisms of experience-dependent neuroplasticity have mostly focused on sensory cortices and how sensory inputs reorganize neuronal circuitry^6^, it remains largely unknown how circuits involved in higher-order cognitive control processes reorganize over the course of learning a task. Such cognitive control tasks require the integration of information streams across multiple brain regions, including the frontal, temporal, and parietal lobes^7^. Within these interconnected regions, previous studies have highlighted the thalamoprefrontal circuit as a conserved substrate for cognitive control across species ^6,8^. Specifically, the mediodorsal thalamic (MdT) projection to the dorsomedial prefrontal cortex (dmPFC), plays an important role in maintaining working memory during task performance, representing current goal values, and reinforcing task-relevant neural activity that give rise to adaptive behaviors ^7,9,10^. In spite of these findings, studies have not assessed whether or how MdT–dmPFC thalamoprefrontal circuitry evolves over the course of learning a task. Such insight could be essential for understanding how the brain dynamically adapts over time to complex cognitive paradigms and provides a novel avenue for exploration in learning and neurodevelopmental disorder models.

To address this issue, we leveraged multi-fiber photometry in combination with genetically encoded calcium indicators to chronically record *in vivo* activity from the mediodorsal thalamic terminals in the dorsomedial prefrontal cortex as mice performed a psychomotor vigilance task over time. During this task, animals learned a sequential association between two tones—one lower frequency (initial cue) followed by one higher (go cue)—to earn a reward. The animals had to learn to respond only after the second tone (go cue) to get the reward. Notably, the inter-trial interval (ITI) varied on a pseudorandom, trial-by-trial basis, preventing mice from knowing the exact timing of initial tone onset. We then studied an analogous paradigm in humans to assess thalamoprefrontal electrical circuit interactions during a homologous task using intracranial electrode recordings. This approach enabled us to capture multimodal thalamoprefrontal dynamics at a millisecond scale, allowing comparison of functional micro and mesocircuit plasticity changes across species.

## Results

### Mouse behavioral paradigm

To assess how the mediodorsal thalamic projections to the dorsomedial prefrontal cortex (MdT-dmPFC circuit) change while learning a psychomotor vigilance task, we trained six mice to perform a type of temporal expectancy task (Figure 1).

**Figure 1:**
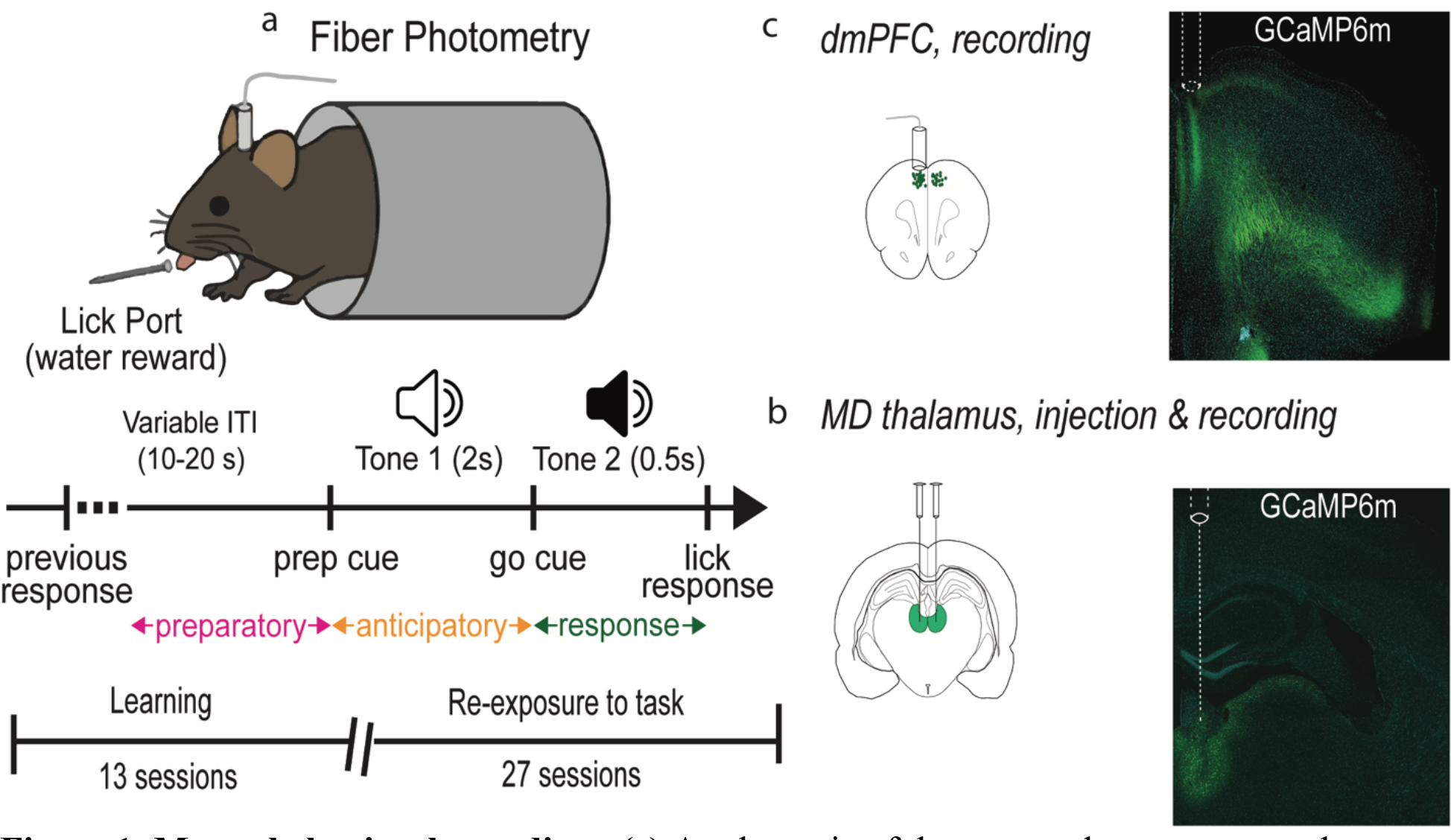
Mouse behavioral paradigm. (a) A schematic of the temporal expectancy task modified for mouse behavior with temporal expectancy cues. (b) Viral injection site into mediodorsal thalamus and (c) fiber placement into the dorsomedial prefrontal cortex. Adapted from Hedlund M, et al. (2025).

For this task, mice were head fixed, and water restricted under protocols approved by our Institutional Animal Care and Use Facility (IACUC protocol #32908). Initially, a low frequency tone was played for two seconds, followed by a high-frequency tone played for half a second. Mice had to respond only after the second tone to receive water as a reward. This behavioral paradigm was based on a previously validated protocol (Lovette-Barron et al., 2017) but was modified to include two tones instead of one to mirror the human version of this psychomotor vigilance task ^11^. This modification enabled a specific distinction between the pre-trial (preparatory) period and the trial–engaged (anticipatory) period.

### Human behavioral paradigm

We employed a homologous task paradigm in 24 human subjects implanted with intracranial electrodes (iEEG) (Figure 2).

**Figure 2:**
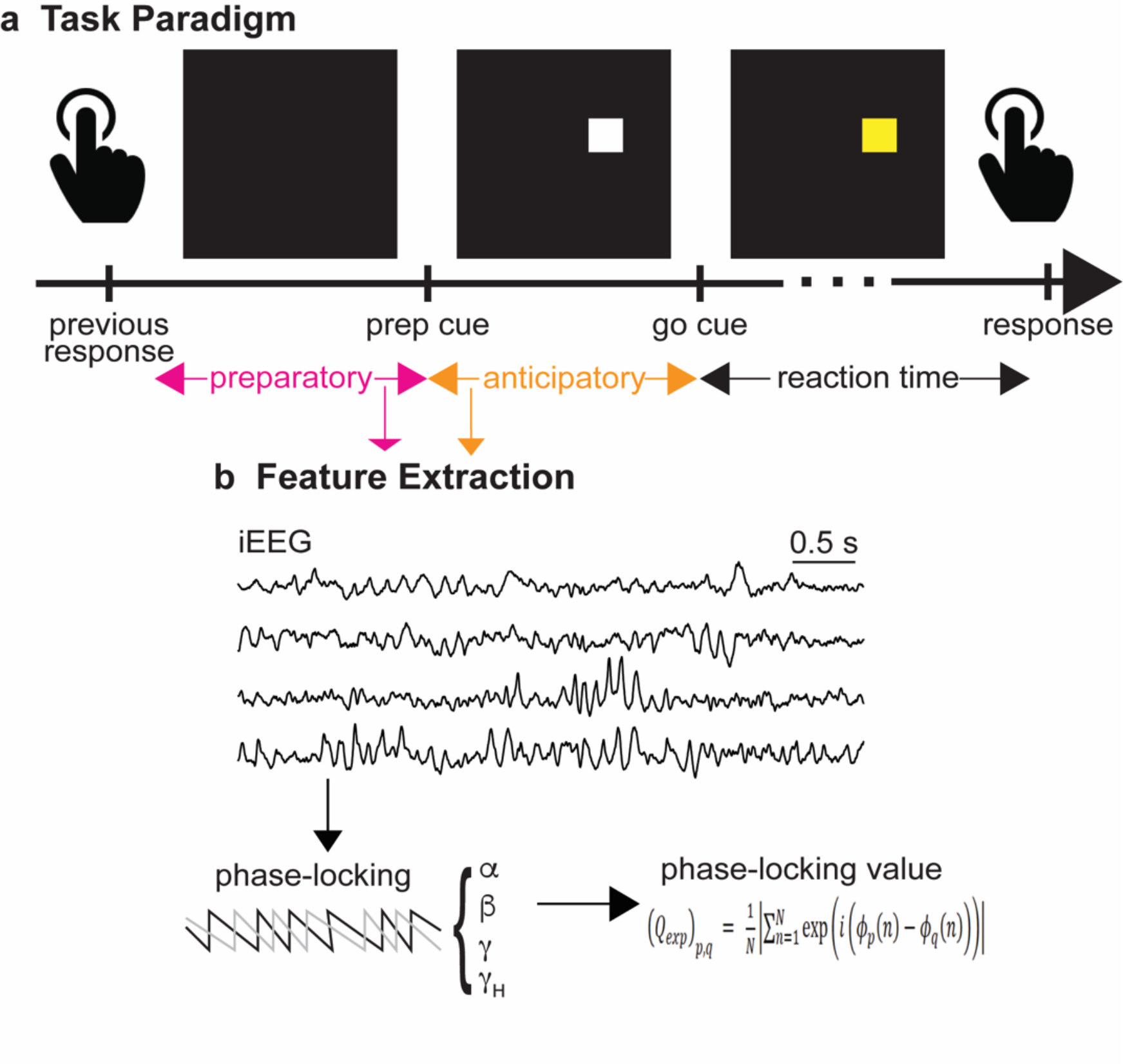
Human behavioral paradigm. (a) A schematic of the temporal expectancy task in humans. (b) Node-feature pairs are quantified from iEEG data and the dynamic brain state was assessed by calculating phase locking between nodes for each of the four canonical frequency bands in the 500ms preparatory, anticipatory, and reward time windows.

Twenty subjects contained the desired thalamoprefrontal coverage and were used for analysis. Dynamic synchrony in circuit nodes was assessed by calculating phase locking between nodes for the theta/alpha and high gamma canonical frequency bands in the 500ms preparatory, anticipatory, and reward retrieval windows. We calculated a time resolved pairwise phase locking value at each time point over early (first third of trials) and late (last third of trials) learning sessions.

### Human and mouse behavior over the course of learning

To assess cognitive task performance across both humans and mice over time, we measured the reaction time (RT) to reward retrieval during the early (first third of trials) and late (last third of trials) learning sessions across species (Figure 3).

**Figure 3:**
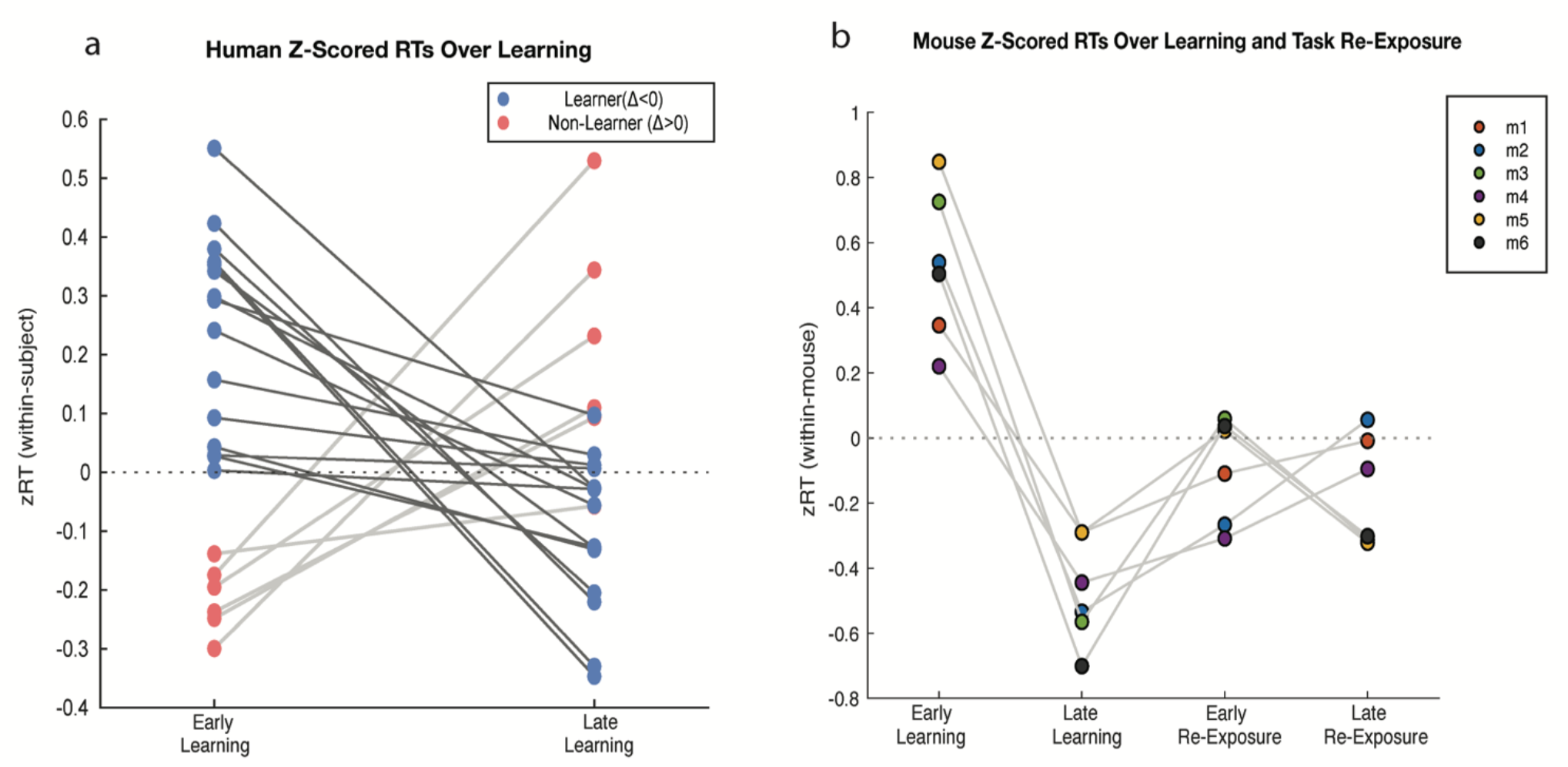
Human and mouse behavior over the course of learning. (a) For humans, reaction times (RTs) from all valid non-error trials (RT< 999 ms) were z-scored within each subject and session. Mean z-scored RTs were computed for the early learning (first third of trials) and late learning (last third of trials) sessions for each subject, which were used to classify Learners (Δ z-scored RT < 0) (blue) and non-Learners (Δ z-scored RT > 0) (red). Lines show paired Early to Late Learning z-scored RT values for each subject. (b) For each mouse and session (learning and re-exposure), RTs from successful trials (RT<500 ms) were z-scored within each mouse and plotted for the Early (first third of trials) and Late (last third of trials) trials of each session — Early Learning, Late Learning, Early Re-exposure, and Late Re-exposure. Dot-and-line plots show per-mouse z-scored RTs across each session, with a unique color assigned to each mouse and light grey lines connecting points across time.

The RTs were normalized to each subject’s mean and standard error of the mean. When assessing RT during learning in humans, we observed two groups, one that acquired the reward faster (decrease in RT) (p<0.001) and one that acquired the reward slower (increase in RT) over time (p<0.001). The group with a decrease in RT was classified as “learners” (n=15), and the group that had an increase in RT was classified as “non-learners” (n=5) (Figure 3a). During the learning sessions in mice, we found that all subjects had a decrease in RT over time (p<0.001). Two weeks following learning, we re-exposed mice to the task and observed a “learner” and “non-learner” group emerge (Figure 3b).

### Learning-induced thalamoprefrontal circuit evolution in mice

To measure activity from the MdT-dmPFC circuit during psychomotor task performance over learning and re-exposure sessions, we targeted a fluorescent Ca2+ indicator (GCAMP6m)-carrying AAV to CAMKIIa+ neurons in the medial dorsal thalamus. We used fiber photometry (Kim et al., 2016) to record changes in fluorescence at the MdT terminals located in the dmPFC, expressing GCAMP6m.

Fiber photometry signals were normalized and visualized as z-scored ΔF/F traces aligned to task events (Figure 4).

**Figure 4:**
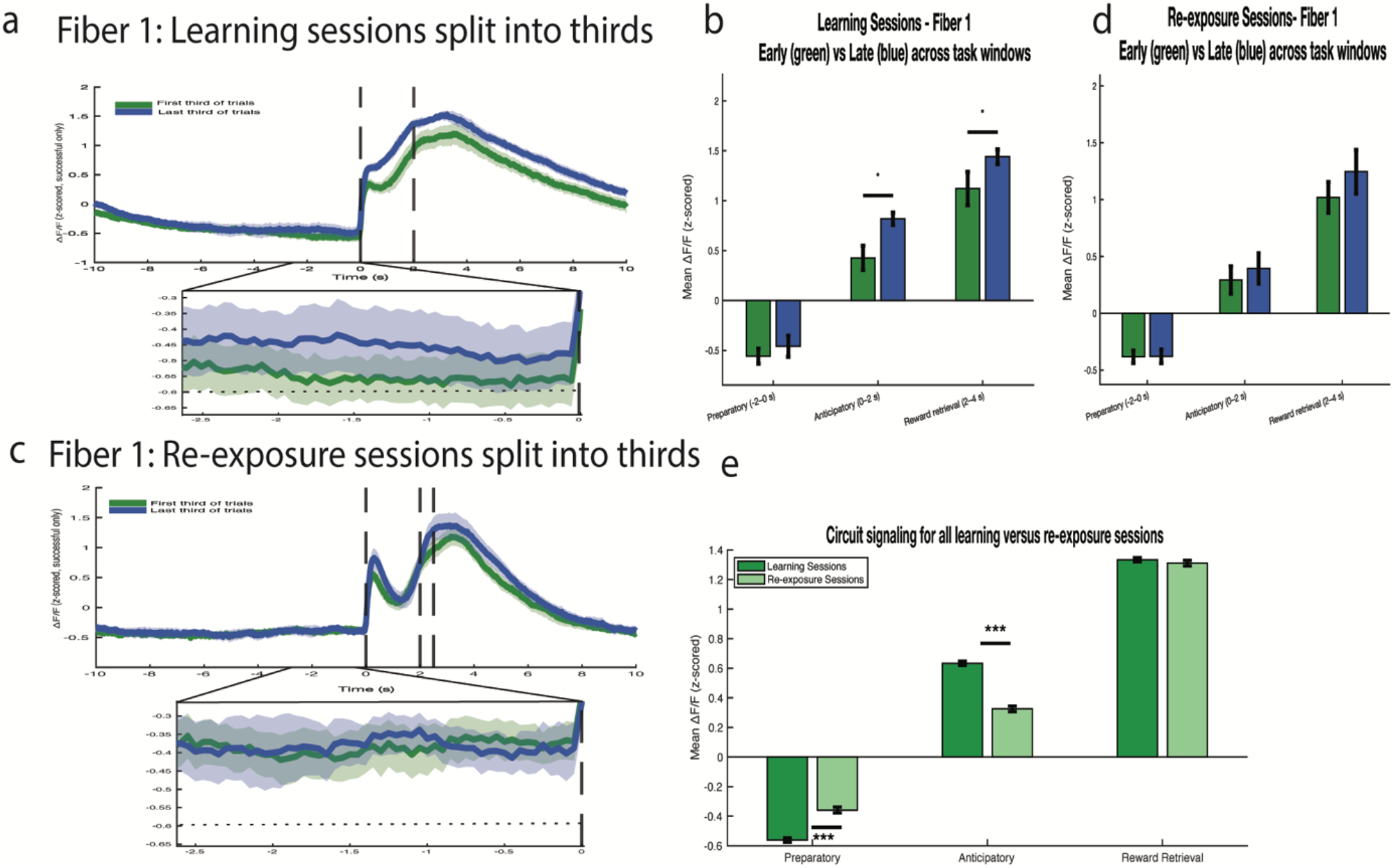
Learning-induced thalamoprefrontal circuit evolution in mice. (a) Fiber photometry ΔF/F traces from successful trials were separated into the first and last third of trials for Learning and (c) Re-Exposure sessions across all mice. Mean z-scored ΔF/F traces with shaded regions indicating the standard error of the mean, are shown aligned to the initial cue (time 0 s). Bar plots quantify the ΔF/F activity in the first and last third of trials for (b) learning and (d) re-exposure sessions. (e) The mean z-scored ΔF/F for Fiber 1 was averaged across all learning and re-exposure trials within the Preparatory, Anticipatory, and Reward Retrieval time windows. Trials were pooled across mice (n=6) and only successful trials with valid reaction times were included (RTs<500ms). Bars reflect the across-trial mean of per-trial window averages; error bars denote the standard error mean across trials. Calcium signaling during learning and re-exposure sessions was assessed within each time window using two-sample *t*-tests with Benjamini–Hochberg FDR corrections (q = 0.05). Asterisks indicate significance (*p* < 0.05, p< 0.01, p < 0.001).

For each recording channel, the raw fluorescence signal was linearly regressed against its corresponding reference channel to correct motion and photobleaching artifacts. The residual fluorescence was then converted to ΔF/F and z-scored using the mean and standard deviation of the entire recording session, yielding normalized activity values expressed in standard deviation units. Event-aligned traces were extracted from −10 s to +10 s relative to tone onset and averaged across trials and animals. The resulting averages represent mean changes in normalized calcium activity across trials, with shaded regions indicating the standard error of the mean. To understand how the fluorescence signal differs over time during learning and re-exposure sessions, we divided the trials into thirds and plotted the first and last third of trial traces (Figure 4a, c). During the learning sessions, we observed that the last third of trials exhibited an increase in calcium signaling prior to the go-cue (anticipatory period) (p<0.05) and reward retrieval period (Figure 4b) (p<0.05). Though we observed no differences between the first and last of trials during the re-exposure sessions (Figure 4d).

To explore whether fluorescent signaling differs between the learning and re-exposure periods, we compared the mean calcium signaling among all learning and re-exposure trials during the preparatory, anticipatory, and reward retrieval time periods. When comparing these two groups, during task re-exposure, we observed an increase of circuit signaling during the preparatory period (p<0.001), a decrease during the anticipatory period (p<0.001) and no difference during reward retrieval (Figure 4e).

### Learning-induced thalamoprefrontal circuit evolution in humans

Having identified a temporal change in thalamo-prefrontal circuit signaling dynamics as mice learned to perform the psychomotor vigilance task, we next asked if similar changes occur at a mesoscale level and whether this is an evolutionary conserved mechanism. To address this question, we analyzed intracranial electrode recordings from twenty human patients performing an analogous temporal expectancy task to measure thalamo-prefrontal connectivity of “learner” and “non-learner” groups. We calculated pairwise time-resolved phase locking values in the high gamma and theta/alpha bands over the first and last third of trials in subjects that contained electrodes in the anterior corona radiata (ACR) and the superior frontal gyrus (SFG) (Figure 5).

**Figure 5:**
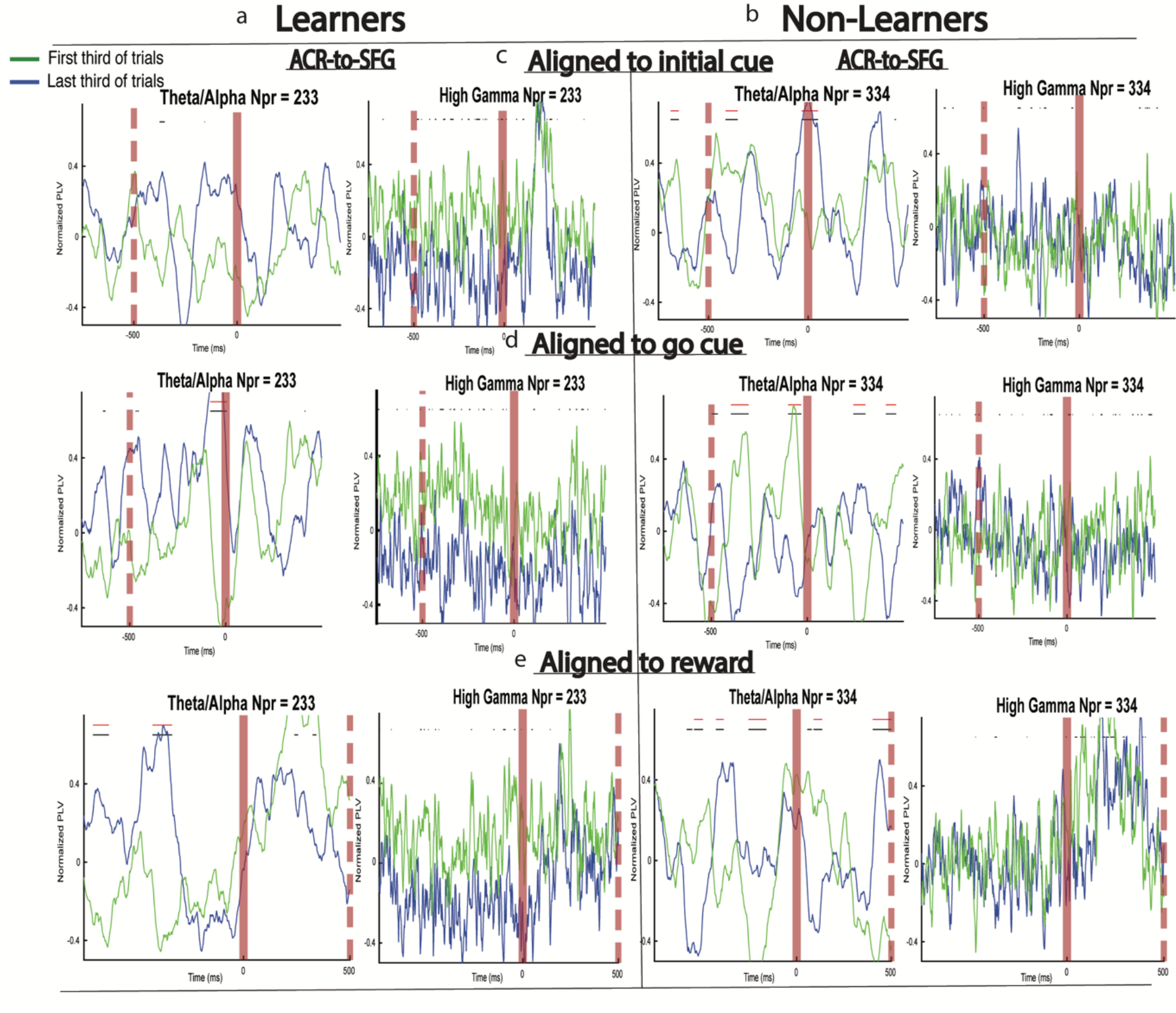
Learning-induced thalamoprefrontal circuit evolution in humans. Time-resolved PLV in thalamoprefrontal pairs for (a) Learners (n=15) and (b) non-Learners (n=5) groups. Time resolved PLV between the anterior corona radiata and the superior frontal gyrus during the (c) preparatory, (d) anticipatory, (e) and reward time periods in the theta/alpha and high gamma frequency bands, calculated over the first third (green) and last third (blue) of trials for Learner and non-Learner groups. In each subplot, PLV is averaged over all pairs for each group between the specified regions across all subjects. The black bar at the top of the plot represents a statistically significant difference between the first and last third of trials, and the red bar denotes the significance of at least twenty consecutive trials.

For the learner and non-learner groups, we observed that the ACR showed significant separation between the early and late sessions in high and low frequency connection to the SFG in the preparatory (500ms prior to initial cue) and anticipatory (500 ms prior to the go cue) time periods (Figure 5a, b).

### Thalamoprefrontal connectivity tilt separates learners from non-learners

To summarize the strength of separation we observed between the first third of trials and the last third of trials between the ACR and the SFG, we analyzed the effect size of change in time-resolved phase locking values in both theta/alpha and high gamma bands in the preparatory, anticipatory, and reward time windows of learners and non-learners (Figure 6).

**Figure 6:**
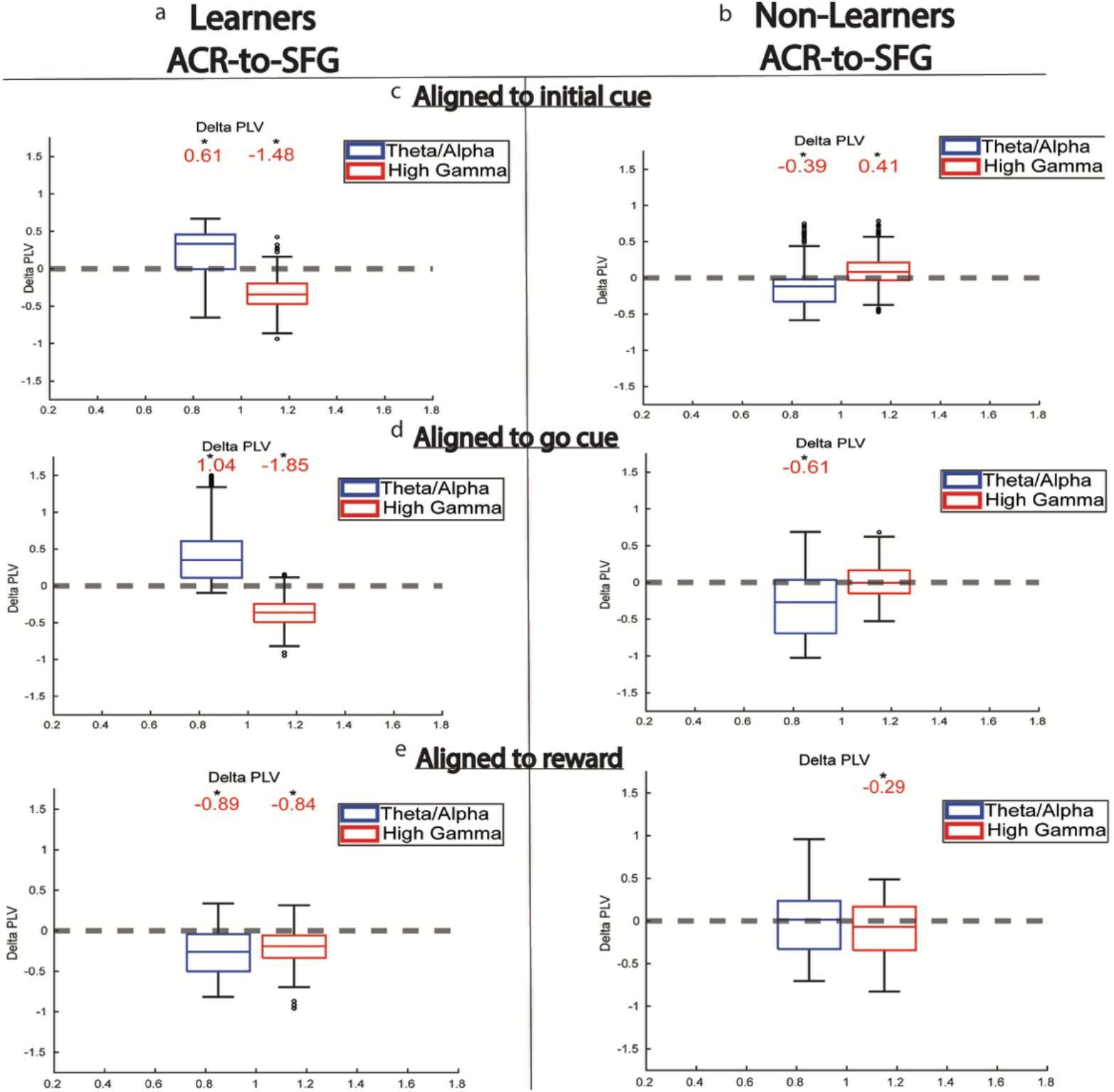
Thalamoprefrontal connectivity tilt separates learners from non-learners. Difference in time-resolved PLV between the first third of trials and last third of trials for the theta/alpha band (blue) and the high gamma band (red) for (a) Learners and (b) Non-Learners. Box plots show the difference at each sample in the (c) preparatory, (d) anticipatory, and (e) reward time windows. The numbers above each box are the Cohen’s d effect size of the sample-by-sample difference between the first and last third of trials for theta/alpha and high gamma. No number means the corresponding Wilcoxon signed-rank test was not significant. Asterisks indicate significance (*p* < 0.05).

We found a strong effect size in the low and high frequency bands in the preparatory and anticipatory time periods for both learners and non-learners. Specifically, during the preparatory period, learners had an increase in theta/alpha connectivity (d=0.61, p<0.05) and a decrease in high gamma connectivity (d=-1.48, p<0.05), as learning sessions progressed. A similar trend was observed during the anticipatory period, where learners had an increase in theta/alpha connectivity (d=1.04, p<0.05) and a decrease in high gamma connectivity (d=-1.85, p<0.05) as learning sessions progressed, demonstrating a connectivity tilt. However, non-learners had an increase in high gamma connectivity (d=0.41, p<0.05) and a decrease in theta/alpha connectivity (d=-0.39, p<0.05) during the preparatory period as learning sessions progressed, demonstrating an opposite connectivity tilt effect. A similar trend was observed during the anticipatory period, with a decrease in theta/alpha connectivity (d=-0.61, p<0.05), as learning sessions progressed. The average difference in effect size across all learners versus non-learners was calculated during the preparatory (2.07), anticipatory (3.5), and reward (0.24) time periods, demonstrating large effect size differences in only the preparatory and anticipatory time windows.

### Conclusion and Outlook

This study presents a novel understanding of how thalamoprefrontal circuit dynamics across mice and humans may reorganize during learning. We found experience-dependent plasticity within the thalamoprefrontal circuit at micro and mesoscales. Microscale synaptic dynamics demonstrate a shift from anticipatory to preparatory activity over the course of learning and re-exposure. At a mesoscale level, task learning alters connectivity between brain regions within the thalamoprefrontal circuit, specifically reliable changes to dynamic synchrony patterns between the anterior corona radiata (ACR) and superior frontal gyrus (SFG). Additionally, a connectivity tilt within the ACR-SFG circuitry is able to distinguish human learners from non-learners. Overall, these findings may provide cross-species evidence of novel, conserved thalamoprefrontal circuit mechanisms of adaptive learning, and may provide a novel circuit target for understanding learning and neurodevelopmental cognitive dysfunction.

## Methods

### Mouse model details

All protocols are approved by Stanford’s Administrative Panel on Laboratory Animal Care. We used 6 wildtype C57BL/6J adult male mice (> P45) for this study (The Jackson Laboratory, #000664). All mice were injected bilaterally with 350 nL of AAV8-CaMKIIa-GCaMP 6m-NRN (8.76 × 10^12^ viral particles/ml). No statistical methods were employed to predetermine sample size, and animal selection was not randomized. Differences between male and female mice were not explored. Mice were singly housed, maintained under a 14:10 reverse light cycle, and had ad libitum access to food and water until the commencement of water restriction (see below). All testing was conducted during the morning and afternoon.

### Mouse behavioral paradigm

The temporal expectancy task includes an auditory presentation cue, a variable delay period, and an auditory go cue of a different pitch after which the mice were trained to elicit a lick response (Figure 1). The first tone was low-frequency 5 kHz, 90 dB, 2 s long, followed by a second high-frequency 20 kHz, 90 dB, 0.5 s long tone, presented at a pseudorandom inter-stimulus interval ranging from 11 to 21 seconds. The preparatory period was defined as approximately 2 s before the onset of the first tone, the anticipatory period was defined as the duration of the first tone (2 s), and the reward retrieval period was defined as 2 s after the second tone. Mice were water restricted for 2 days prior to beginning training and were maintained at > 80% of their pre-deprivation weight throughout training. Mice were habituated to head fixation and were trained to obtain a water reward (7 µL) triggered by frequency changes in auditory stimuli by contacting a lick port with their tongues over the course of 2-3 weeks. During this time, a water reward was not given if mice licked in response to the first tone. Mice were head-fixed in a plastic tube using custom-designed head plates (Protolabs) implanted during surgery (see below). Behavioral protocols were implemented and run on a Bpod State Machine r2 (Sanworks) using a proprietary graphical user interface that communicated with MATLAB (MathWorks). Behavioral responses were recorded using an optical lickometer (Sanworks). Sounds were delivered using a piezoelectric loudspeaker (Diginex) positioned ~35 cm from the mouse’s head and were generated by the Bpod State Machine r2 (Sanworks). The mice were placed in an acoustically isolated behavior chamber (Med Associates) which was illuminated using a mounted infrared LED (ThorLabs). The behavior was recorded using a high-speed camera (Basler) with a 24 mm/F1.4 lens (Edmund Optics) focused on the tongue.

### Mouse surgery

Mice were anesthetized with 4% isoflurane and maintained with 2%–3% isoflurane in a stereotactic frame. Following stereotaxic affixation, ophthalmic ointment was applied to prevent eye dryness. The head was cleaned with betadine antiseptic solution (Betadine) and 70% isopropanol wipes. The scalp and periosteum were removed, and the skull cleaned thoroughly with 3% hydrogen peroxide solution and saline. A small craniotomy was performed over the injection site (MD thalamus). The virus (250–500 nL) was injected with a 10 μL syringe and a beveled metal needle (World Precision Instruments) at a rate of 150 nL/min, controlled by a pump controller connected to an injection pump (World Precision Instruments). The syringe was slowly withdrawn 25 minutes after the injection. Following the virus injection, a 400 μm optic fiber (Doric) was slowly lowered to within 50–100 μm of the target site and fixed in place using tissue adhesive (3M Vetbond) and adhesive cement (C&B Metabond, Parkell) along with a custom aluminum headplate. For photometry experiments, two mono fiber-optic cannulas (Doric, MFC_400/430-0.48_4/2mm_MF1.25_FLT) were placed unilaterally above the MD and dmPFC, with alternating hemispheres. Mice received 0.05 mg/kg Buprenorphine for analgesia and were placed on a warming pad to recover from anesthesia. After regaining coordinated locomotion, mice were returned to a clean home cage and monitored daily for the subsequent week. Water restriction began four to five weeks after surgery. The injection coordinates were −1.05 A/P, ±0.5 M/L, −3.35 D/V (MD thalamus), and the fiber implant coordinates were +2.25 A/P, ±0.35 M/L, −1.2 D/V (dmPFC).

### Mouse fiber photometry

Fiber photometry in combination with genetically encoded calcium indicators (GECIs) allows monitoring of cell-and circuit-specific population activity in vivo ^12^. The behavioral setup was based on the protocol described in Lovett-Barron et al. (2017)^13^, and the fiber photometry setup was based on the protocol described by Kim et al. (2016)^14^. Briefly, a custom bundle branching fiber-optic patch cord (Doric) was coupled to two implanted fibers in the MD thalamus and the dmPFC (Doric, 1.25 mm ferrule diameter, 400 μm core diameter, 0.48 NA) for simultaneous activity recording of MD thalamic cell bodies and axons in the dmPFC receiving input from the MD thalamus. Custom MATLAB (Mathworks) routines controlled the timing of the excitation light source, synchronously acquired camera frames, and computed the total fluorescence from each fiber in real-time. Artifact subtraction was performed using the 405 nm reference channel. For quantification and comparison across days and animals, all data was z-scored with baseline defined as the mean of the entire reference-subtracted photometry signal for each trial.

### Mouse analyses

Mouse behavior was determined from the licks recorded on the lickometer and the timing of auditory stimulus onset. Reaction times (RTs) were defined as the time to the first lick after the onset of the second tone, within its duration (500 ms). Trials were counted as successful if mice initiated licks within the duration of the second tone, with no licks during or within 10 s before the first tone. For each type of experiment, the first trial of each daily session was excluded to mitigate the effects of lickometer location adjustment at the beginning of the session. Each photometry experiment dataset comprised one hour of recording per mouse.

### Mouse immunohistochemistry

Mice were deeply anesthetized and injected with 0.1 mL of EUTHASOL^®^ (pentobarbital sodium and phenytoin sodium) euthanasia solution (Virbac Corporate). They were then transcardially perfused with ice-cold PBS, followed by 4% PFA in PBS. Brains were removed and placed in 4% PFA in PBS for post-fixation overnight at 4°C on a shaker. The brains were then mounted and sectioned into 150 μm coronal sections using a vibrating blade microtome (Leica). Sections were placed into PBS in 24-well plates, washed at room temperature in PBST, then incubated with GFP Polyclonal Antibody, AF488, Cat. # A21311 (1:350) (ThermoFisher Scientific) to amplify GCaMP signal. DAPI was added prior to mounting (1:500) (Invitrogen). Sections were subsequently mounted on slides using mounting media and imaged with confocal microscopy (Olympus Life Science).

### Human subject details

We studied 20 human subjects (aged 18-56; 15 females) who were implanted with bilateral iEEG as part of a clinical evaluation for epilepsy surgery and consented to research performed under approval of the Institutional Review Board of the University of Pennsylvania. Two subjects had below average intelligence as measured by the Wechsler Adult Intelligence Scale (WAIS-IV) test^15^. We used AdTech depth electrodes.

### Human behavioral paradigm

All subjects performed a temporal expectancy task, with three subjects completing two sessions of the task. Each task trial consisted of a visual preparatory cue, a variable delay period of either 500 or 1500 ms, and a visual go cue after which each subject must press a key in response (Figure 2). The inter-trial interval was 2500-2750ms following the response. To study connectivity, we extracted features from the preparatory period (500 ms prior to the presentation cue for each trial), task-engaged anticipatory period (500 ms prior to the go cue for each trial), and reward phase (500 ms following the go cue for each trial).

### Human time-resolved PLV

Time-resolved PLV (trPLV) was used to analyze pairwise thalamo-prefrontal interactions averaged over trials. TrPLV was calculated between each pair of electrodes over a set of trials for each sample in a time window of interest. For every subject and task session, this was done for the first third of trials and last third of trials. For each time window of interest (time = 0 aligned to initial cue, go cue, or reward), the baseline period was defined as −750 to −500 ms. To compare PLV calculated across the first third of trials and PLV calculated across the last third of trials, the baseline for both first third and last third trial PLVs was averaged for each pair, and pairwise PLV was normalized to this averaged baseline. Pairwise trPLV was averaged over all pairs between the anterior corona radiata and the superior frontal gyrus across all subjects and sessions. We used a two-sample Wilcoxon signed-rank test between the averaged first third of trial PLVs and average last third of trials PLVs at each time point to quantify the difference in phase-locking behavior between each set of trials. Bonferroni corrections were used to adjust the p-value significance level to 1.96e-5 to account for 1278 multiple comparisons (2 frequency bands times 639 samples in the time window of interest). The outcome of this test is depicted in the horizontal black lines at the top of each subplot, and a red horizontal line depicts where a significant difference exists over at least 20 continuous samples (Figure 5). Cohen’s d effect size was used to quantify the strength of difference between the average first third of trials and average last third of trials PLV over all samples in the preparatory, anticipatory, and reward periods (Figure 5).

### Human connectivity tilt

The connectivity tilt phenomenon is characterized by a behaviorally relevant increase in low frequency pairwise synchrony and concomitant decrease in high frequency pairwise synchrony. We quantified connectivity tilt by analyzing the strength of difference in trPLV between the first third and last third of trials for both low and high frequency interactions. The difference in trPLV for first third and last third of trials is calculated at every time point in the preparatory, anticipatory, and reward time periods. A Wilcoxon sign-rank test, with Bonferroni correction for 3 comparisons (ACR to SFG in 2 frequency bands), is used to determine if the mean difference is significantly different from 0. The Cohen’s d effect size of significant differences quantifies the strength of these effects. Regions that demonstrate a connectivity tilt are those with both a moderate to strong positive effect (d > 0.4) in the theta/alpha band and a moderate to strong negative effect (d < −0.4) in the high gamma band (Figure 6).

### Statistical analysis

Statistical analyses were performed in MATLAB R2023a (MathWorks). Between-group comparisons of ΔF/F fluorescence used two-sided unpaired *t*-tests, while within-trial epoch comparisons used paired *t*-tests and the false discovery rate (FDR) method for multiple comparisons. Time-resolved PLV analyses were assessed using Wilcoxon signed-rank tests across time points. Cohen’s d effect size was used to quantify strength of statistically significant tests.

## References

1. Kolb B, Mychasiuk R, Muhammad A, Li Y, Frost DO, Gibb R. Experience and the developing prefrontal cortex. Proc Natl Acad Sci U S A. 2012 Oct 16;109 Suppl 2(Suppl 2):17186–93. doi: 10.1073/pnas.1121251109. Epub 2012 Oct 8. PMID: 23045653; PMCID: PMC3477383.

2. Pascual-Leone, A., Amedi, A., Fregni, F., & Merabet, L. B. (2005). The Plastic Human Brain Cortex. Annual Review of Neuroscience, 28(1), 377–401.

3. Wang, M., & Yu, X. (2023). Experience-dependent structural plasticity of pyramidal neurons in the developing sensory cortices. Current Opinion in Neurobiology, 81, 102724.

4. Citri, A., & Malenka, R. C. (2007a). Synaptic plasticity: Multiple forms, functions, and mechanisms. Neuropsychopharmacology, 33(1), 18–41.

5. Hooks, B. M., & Chen, C. (2020). Circuitry underlying experience-dependent plasticity in the mouse visual system. Neuron, 106(1), 21–36.

6. Wang L, Caras ML, Karayannis T, Froemke RC. Editorial: Mechanisms Underlying Experience-Dependent Plasticity of Cortical Circuits. Front Cell Neurosci. 2021 Apr 30;15:687297. doi: 10.3389/fncel.2021.687297. PMID: 33994953; PMCID: PMC8121167.

7. Jung J, Cloutman LL, Binney RJ, Lambon Ralph MA. The structural connectivity of higher order association cortices reflects human functional brain networks. Cortex. 2017 Dec;97:221–239. doi: 10.1016/j.cortex.2016.08.011. Epub 2016 Sep 2. PMID: 27692846; PMCID: PMC5726605.

8. Sébastien Parnaudeau, Scott S. Bolkan, Christoph Kellendonk. The Mediodorsal Thalamus: An Essential Partner of the Prefrontal Cortex for Cognition, Biological Psychiatry, Volume 83, Issue 8, (2018).

9. Miller, R. L. A., Francoeur, M. J., Gibson, B. M., & Mair, R. G (2017). Mediodorsal thalamic neurons mirror the activity of medial prefrontal neurons responding to movement and reinforcement during a dynamic DNMTP Task. Eneuro, 4(5).

10. Bolkan, S., Stujenske, J., Parnaudeau, S. et al. Thalamic projections sustain prefrontal activity during working memory maintenance. Nat Neurosci 20, 987–996 (2017)

11. Buch, V. P., Bernabei, J. M., Ng, G., Richardson, A. G., Ramayya, A., Brandon, C., Stiso, J., Bassett, D. S., & Lucas, T. H. (2022). “Primed to Perform:” Dynamic White Matter Graph Communicability May Drive Metastable Network Representations of Enhanced Preparatory Cognitive Control.

12. Patel, A. A., McAlinden, N., Mathieson, K. & Sakata, S. Simultaneous Electrophysiology and Fiber Photometry in Freely Behaving Mice. Front Neurosci 14, 148 (2020).

13. Lovett-Barron, M. et al. Ancestral Circuits for the Coordinated Modulation of Brain State. Cell 171, 1411–1423 e1417 (2017).

14. Kim, C. K. et al. Simultaneous fast measurement of circuit dynamics at multiple sites across the mammalian brain. Nat Methods 13, 325–328 (2016).

15. Wechsler, D. WAIS-IV Administration and Scoring Manual (Wechsler Adult Intelligence Scale—Fourth Edition). (NCS Pearson, 2010).

16. Alcaraz F, Fresno V, Marchand AR, Kremer EJ, Coutureau E, Wolff M. Thalamocortical and corticothalamic pathways differentially contribute to goal-directed behaviors in the rat. Elife. 2018 Feb 6;7:e32517. doi: 10.7554/eLife.32517. PMID: 29405119; PMCID: PMC5800843.

17. DeNicola, A. L., Park, M.-Y., Crowe, D. A., MacDonald, A. W., & Chafee, M. V. (2020a). Differential roles of mediodorsal nucleus of the thalamus and prefrontal cortex in decision-making and state representation in a cognitive control task measuring deficits in schizophrenia. The Journal of Neuroscience, 40(8), 1650–1667.

18. Buch, V.P., Brandon, C., Ramayya, A.G. et al. Dichotomous frequency-dependent phase synchrony in the sensorimotor network characterizes simplistic movement. Sci Rep 14, 11933 (2024).

